# White matter connectivity of uncinate fasciculus and inferior fronto-occipital fasciculus: A possible early biomarker for callous-unemotional behaviors in young children with ADHD

**DOI:** 10.1101/2020.08.19.257691

**Authors:** Paulo A. Graziano, Dea Garic, Anthony S. Dick

**Author notes:** Corresponding Author Information: Paulo Graziano, Ph.D., Center for Children and Families & Department of Psychology Florida International University, 11200 SW 8th Street, AHC 4 Rm. 459, Miami, Florida 33199.

## Abstract

**Background:** Callous-unemotional (CU) behaviors are important for identifying severe patterns of conduct problems (CP). One major fiber tract implicated in the development of CP is the uncinate fasciculus (UF), which connects amygdala and orbitofrontal cortex (OFC). The goals of the current study were to 1) explore differences in the white matter microstructure in the UF and other major fiber tracks (inferior longitudinal fasciculus [ILF], inferior fronto-occipital fasciculus [IFOF], cingulum, and corticospinal tract [CST]) between *young* typically developing (TD) children and those with Attention-Deficit/Hyperactivity Disorder (ADHD), and 2) explore, within the ADHD group, whether individual differences in these white matter microstructures relate to co-occurring CP and CU behaviors, respectively.

**Methods:** Participants included 198 young children (78% boys, *M*_age_ = 4.95 years; 80% Latinx; 49% TD). CU behaviors and CP were measured via a combination of teacher/parent ratings. Non-invasive diffusion-weighted imaging (DWI) was used to measure fractional anisotropy (FA), an indirect indicator of white matter properties.

**Results:** Relative to TD children, children with ADHD had reduced FA on four out of the five fiber tracks we examined (except for cingulum). Within the ADHD group, no associations were found between CP and reduced white matter integrity across any of the fiber tracks examined. However, we found that even after accounting for CP and a host of covariates including whole brain FA, CU behaviors were independently related to reduced FA in bilateral UF and left IFOF.

**Conclusions:** The bilateral UF and IFOF may be a biomarker of CU behaviors, even in very young children.

---

Young children exhibiting early signs of conduct problems (CP), typically represented by disruptive behavior disorder diagnoses such as attention-deficit/hyperactivity disorder (ADHD), oppositional defiant disorder (ODD), and/or conduct disorder (CD), represent the most common referrals to mental health clinics (2013; Polanczyk, Willcutt, Salum, Kieling, & Rohde, 2014). A significant factor identified as contributing to the heterogeneity present in the manifestation of early CP is callous-unemotional (CU) traits, which refer to low levels of guilt, empathy, and caring for others (Frick, Ray, Thornton, & Kahn, 2014). CU *traits* or *behaviors*^*\**^, a more developmentally appropriate term to refer to the CU construct in early childhood, can be reliability identified in the preschool period (Ezpeleta, de la Osa, Granero, Penelo, & Domenech, 2013; Waller, Hyde, Grabell, Alves, & Olson, 2015) and these have been an important construct for identifying the most pervasive, severe, and aggressive patterns of CP and later antisocial behavior (Frick et al., 2014). Not surprisingly, emerging neuroscience studies have started to examine the neural signatures, both at the structural and functional level, with the current study focusing on the potential disrupted *connectivity* between brain regions as a way to understand the development of CP and/or CU behaviors.

## Connectomic Differences Associated with CP/CU

The fiber pathways comprising the structural connectome among extended limbic, frontal, and temporal regions have been the main subject of inquiry as it relates to CP/CU. Diffusion-weighted imaging (DWI), a non-invasive MRI technique that measures the diffusion of water molecules along anisotropic fiber bundles (Beaulieu, 2002), has been the method of choice for investigating the structural network of fiber pathways. Most studies have focused on differences in fractional anisotropy (FA) (Winston, 2012). Higher FA values index a greater anisotropic (directional) water diffusion within axonal fibers, which is taken as a general index of fiber integrity (Soares, Marques, Alves, & Sousa, 2013; Thomason & Thompson, 2011).

Using this technique, researchers have attempted to determine whether disrupted connectivity between brain regions is associated with the development of CP and CU behaviors (see Waller, Dotterer, Murray, Maxwell, and Hyde (2017) for review). For example, some researchers have suggested that disrupted connectivity between amygdala and prefrontal cortex is associated with the development of CP and CU behaviors (Blair, 2007), contributing specifically to the underlying cognitive, reward, and emotional processing mechanisms related to CP/CU (Raine, 2018). Fronto-amygdala connectivity is accomplished in part via the uncinate fasciculus (UF). This fiber pathway has rostral terminations in orbital and lateral frontal cortex, frontal pole, and anterior cingulate gyrus. The posterior termination in the temporal lobe includes projections through amygdala (de Schotten, Dell’Acqua, Valabregue, & Catani, 2012; Holl et al., 2011; Von Der Heide, Skipper, Klobusicky, & Olson, 2013).

Several studies have found reduced FA in the UF among adult samples exhibiting high levels of CP (Craig et al., 2009; Motzkin, Newman, Kiehl, & Koenigs, 2011; Sobhani, Baker, Martins, Tuvblad, & Aziz-Zadeh, 2015). The only studies of youth have been conducted in adolescents (Waller et al., 2017). In these cases, reduced FA in UF is associated with increased CU behaviors (Breeden, Cardinale, Lozier, VanMeter, & Marsh, 2015) and increased psychopathy (Maurer, Paul, Anderson, Nyalakanti, & Kiehl, 2020), although Sarkar and colleagues (2013) and Passamonti and colleagues (2012) reported the opposite association. Other abnormalities of the fiber pathways supporting extended limbic, frontal, and temporal regions have also been reported (Waller et al., 2017). In particular, fiber pathways of the ventral temporal lobe, namely the inferior longitudinal fasciculus (ILF) and inferior fronto-occipital fasciculus (IFOF), have been associated with psychopathic traits and conduct disorder in adolescents (Haney-Caron, Caprihan, & Stevens, 2014; Pape et al., 2015). The ILF courses in the ventral white matter of the temporal lobe, originating posteriorly in extrastriate areas of the occipital lobe, and ending with rostral terminations in the middle and inferior temporal gyri, the temporal pole, parahippocampal gyrus, hippocampus, and amygdala (Catani, Jones, Donato, & Ffytche, 2003). The IFOF runs medial to the ILF, originates in the inferior and medial occipital lobe, travels through the temporal stem dorsal to the UF, and projects to the inferior frontal gyrus, the medial and orbital frontal cortex, and the frontal pole (Catani et al., 2003; Martino, Brogna, Robles, Vergani, & Duffau, 2010; Martino, Vergani, Robles, & Duffau, 2010; Sarubbo, De Benedictis, Maldonado, Basso, & Duffau, 2013). These two pathways connect a number of limbic, frontal, and temporal regions associated with CP/CU, and thus these findings are predictable in that context. Finally, mixed findings in adolescents have been reported for the cingulum (Waller et al., 2017), which is a collection of smaller short association fiber systems that course in the white matter under the cingulate gyrus, supporting connections to/from lateral and dorsal prefrontal cortex, medial prefrontal and anterior cingulate, insula, parahippocampal gyrus, subiculum, and amygdala. The structure and function of these regions, especially insula, amygdala, and anterior cingulate, have been associated with CP/CU. However, only two studies have reported any association in adolescents (2014) (2015).

Although these are promising findings, the literature remains inadequate for understanding the development of CP/CU in very young children. In addition, measurement issues limit the degree to which firm conclusions can be made. For example, often comorbid ADHD is not measured, and therefore it is unclear whether the findings are really due to unmeasured ADHD symptomology (Waller et al., 2017). More focused dissociation of CP with and without high levels of CU behaviors is also needed. Thus, to further our understanding of the neurobiology of CP, more pediatric connectivity studies are needed that take into account CP, CU behaviors, and high comorbidity of ADHD.

### Goals of the Current Study

The overarching goal of the current study was to examine the white matter microstructure in the UF along with other major fiber tracks (ILF, IFOF, and cingulum; see Figure 1) among young typically developing (TD) children and those diagnosed with ADHD. Our goals were to 1) explore differences in these white matter connections between young TD children and those with ADHD, and 2) explore, within the ADHD group evidencing sufficient variability in CP and CU behaviors, whether individual differences in these white matter microstructures relate to co-occurring CP and CU behaviors, respectively. Based on prior work with older youth/adults (Breeden et al., 2015; Waller et al., 2017), we expected children in the ADHD group, which was expected to have high comorbidity rates of CP via ODD/CD diagnoses, to have lower integrity of white matter microstructure across the examined fiber pathways. More specificity in white matter disruption was expected when examining only the ADHD group, as we expected reduced white matter integrity in the UF to be associated with CU behaviors, above and beyond CP.

**Figure 1.**
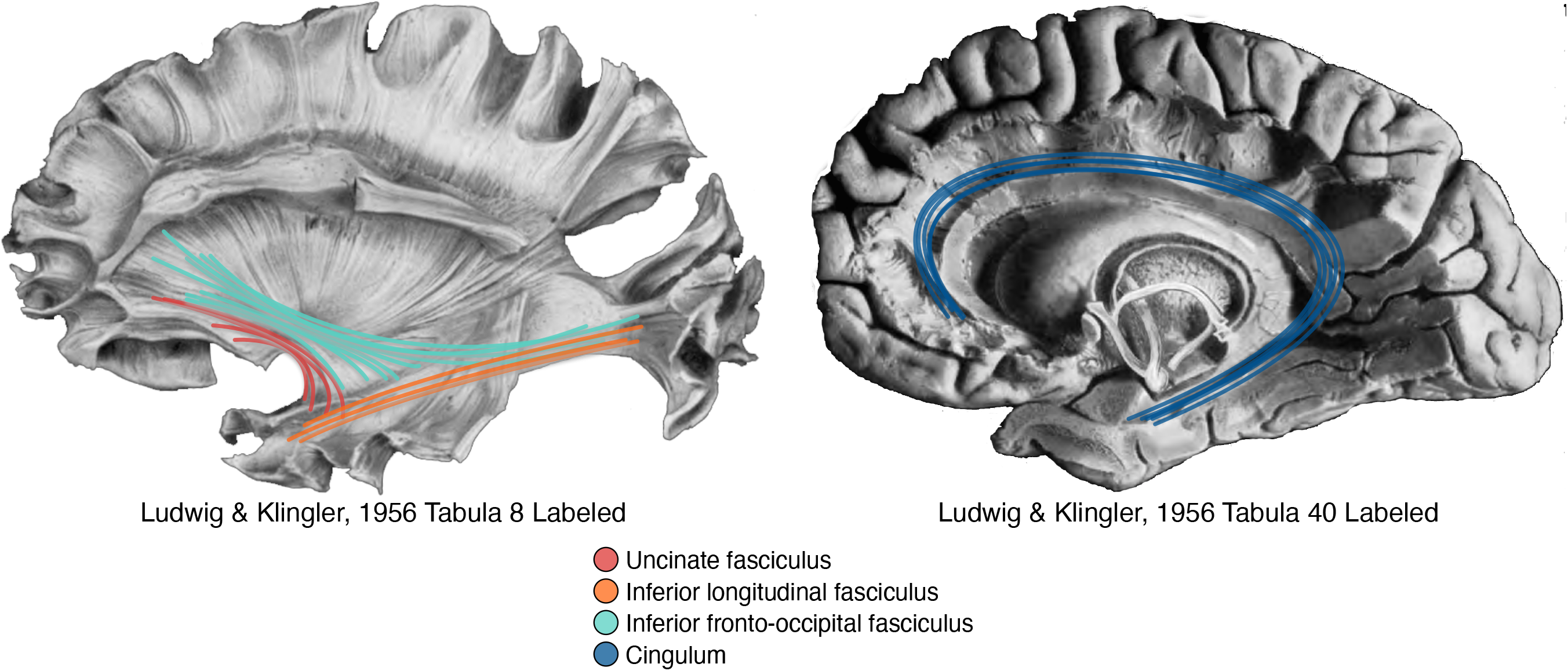
The four fiber pathways of interest are shown overlaid on fiber dissection tabula from Ludwig, E., & Klingler, J. (1956). *Atlas cerebri humani*. Boston and Toronto: Little, Brown, and Company.

## Method

### Participants and Recruitment

The study took place in a large urban southeastern city in the U.S. with a large Latinx population. Children and their caregivers were recruited from local preschools and mental health agencies via brochures, radio and newspaper ads, and open houses/parent workshops. For the ADHD sample, parent and child were invited to participate in an assessment to determine study eligibility if the parent (1) endorsed clinically significant levels of ADHD symptoms, (2) indicated that the child is currently displaying clinically significant academic, behavioral, or social impairments as measured by a score of three or higher on a seven-point impairment rating scale (Fabiano et al., 2006), and (3) were not taking any psychotropic medication. For the TD sample, if the parent (1) endorsed less than 4 ADHD symptoms (across either Inattention or Hyperactivity/Impulsivity according to the DSM-5), (2) less than 4 ODD symptoms, and (3) indicated no clinically significant impairment, the parent and child were invited to participate in an assessment to determine study eligibility. Participants were also required to be enrolled in school during the previous year, have an estimated IQ of 70 based on the WPPSI-IV (Wechsler, 2012), have no confirmed history of an Autism Spectrum Disorder, and be able to attend an 8-week summer treatment program (STP-PreK; Graziano, Slavec, Hart, Garcia, and Pelham (2014) prior to the start of the next school year (ADHD group only).

ADHD diagnosis and comorbid disruptive behavior disorders were assessed through a combination of parent structured interview (Computerized-Diagnostic Interview Schedule for Children [C-DISC]; (Shaffer, Fisher, Lucas, Dulcan, & Schwab-Stone, 2000) and parent and teacher ratings of symptoms and impairment (Disruptive Behavior Disorders Rating Scale, Impairment Rating Scale; (Fabiano et al., 2006; Pelham, Gnagy, Greenslade, & Milich, 1992), as is recommended practice. Dual Ph.D. level clinician review was used to determine diagnosis and eligibility.

The final participating sample consisted of 198 young children (*M*_age_ = 5.70, *SD* = 0.87, and 69% male; 49% TD). Eighty percent of the children were identified by parents as Hispanic/Latino White, 12% as Non-Hispanic/Latino White, 6% as Non-Hispanic/Latino Black, and 2% as Hispanic/Latino Black. The socioeconomic status was low- to middle-class (*M =* $48,477; *SE=* $4,911). Maternal education was input as a binned variable to measure socioeconomic status (SES; 1= some high school thru 6= graduate school/advanced degree; *M =* 4.65; *SE*= 0.19). Of the whole sample, 48.5% were TD (*n* = 96) while the remaining 51.5% met diagnostic criteria for ADHD (*n* = 102). In terms of co-morbidity, 61.76% of children in the ADHD group also met diagnostic criteria for ODD/CD (*n* = 63).

### Study Design and Procedure

This study was approved by the university’s Institutional Review Board. As part of the baseline assessment, children completed a series of tasks in the laboratory and participated in an MRI scanning session. Parents also completed various questionnaires regarding their children’s emotional, behavioral, and cognitive functioning. Families of children with ADHD received the intervention (STP-PreK) at either no cost via a federal grant or at a subsidized cost via a local grant, and all families received compensation ($100 gift card for completing the assessment). Similar questionnaires were also obtained from children’s school teachers. TD children received a $100 gift card, academic and intellectual functioning feedback, study t-shirt, and a small gift from the study “treasure chest.”

### Measures

#### CP

Parents and teachers completed the *Disruptive Behavior Disorders Rating Scale* (DBD; (Pelham et al., 1992), adapted for DSM-5 terminology, which assess for symptoms of ADHD, ODD, and CD on a four point scale with respect to the frequency of occurrence. For the purposes of this study, we obtained an average score for the ODD and CD symptoms (α’s = .71 - .88) as a measure of CP, given their significant correlations (*r*s = .51 - .65, *ps* <.001). Consistent with prior work using the “and/or” algorithm (Piacentini, Cohen, & Cohen, 1992), the highest score among parent and teacher reports was used. To control for ADHD symptom severity, we also examined the hyperactivity/impulsivity and inattention symptoms.

#### Callous-unemotional (CU) Behaviors

Parents (α= .83) and teachers (α= .72) completed a 12 item abbreviated version of the *Inventory of Callous-Unemotional Traits* (ICU; (Frick, 2004; Hawes et al., 2014). An overall CU composite was created by averaging these 12 items and the highest score among parent and teacher reports was used.

##### MRI Acquisition & Processing

All imaging was performed using a research-dedicated 3 Tesla Siemens MAGNETOM Prisma MRI scanner (V11C) with a 32-channel coil located on the university campus. Children first completed a preparatory phase using a realistic mock scanner. In the magnet children watched a child-friendly movie of their choice. Ear protection was used, and sound was presented through MRI compatible headphones.

We collected multi-shell diffusion-weighted imaging data according to the Adolescent Brain and Cognitive Development (ABCD) protocol (Hagler et al., 2019). These scans were collected with a 1.7 mm isotropic voxel size, using multiband imaging echo planar imaging (EPI; acceleration factor = 3). The acquisition consisted of ninety-six diffusion directions, seven b=0 frames, and four b-values (6 b=500, 15 b=1000, 15 b=2000, and 60 b=3000).

##### Diffusion-weighted Imaging Post-Processing

Initial post-processing was accomplished with DTIPrep v1.2.8 (Oguz et al., 2014), TORTOISE DIFFPREP v3.1.0 (Irfanoglu, Nayak, Jenkins, & Pierpaoli, 2017; Pierpaoli et al., 2010), FSL v6.0.1 topup (Andersson, Skare, & Ashburner, 2003; Smith et al., 2004), and DSI Studio (v. June 2020 (Yeh, Wedeen, & Tseng, 2010). We also implemented a pre- and post-analysis quality check assessing signal-to-noise of each diffusion b-value (2016).

Initial quality control was accomplished in DTIPrep to complete the following steps: 1) image/diffusion information check; 2) padding/cropping of data; 3) Rician noise removal; 4) slice-wise, interlace-wise, and gradient-wise intensity and motion checking. The number of acquisitions removed was used as a proxy for movement/bad data quality, and was included as a covariate in subsequent regression analyses. TORTOISE DIFFPREP was used to accomplish motion and eddy current correction. We implemented calculation of the diffusion tensor model in DSI Studio to estimate the eigenvalues reflecting diffusion parallel and perpendicular to each of the fibers along 3 axes (x, y, z). The resulting eigenvalues were then used to compute indices of fractional anisotropy (FA), radial diffusivity (RD), and axial diffusivity (AD; (Basser, Mattiello, & LeBihan, 1994; Hasan & Narayana, 2006). FA is an index for the amount of diffusion asymmetry within a voxel, normalized to take values from 0 (isotropic diffusion) to 1 (anisotropic diffusion). This value can be decomposed into AD, measuring the parallel eigenvalue (λ1), and RD, measuring the average of the secondary and tertiary perpendicular eigenvalues ([λ2+ λ3]/2). AD and RD quantifications are sensitive to axon integrity and myelin integrity, respectively (Basser et al., 1994; Winston, 2012).

##### Fiber Tract Identification

Tractography was conducted using DSI Studio’s built-in tractography atlas (Yeh, 2017). The atlas was originally created from 840 healthy adults in the HCP840 dataset, and defines white matter regions of interest (ROIs) in the MNI space. Because we are analyzing a pediatric dataset, each ROI was visually inspected to ensure that any warping to the atlas template did not introduce inaccuracies. The following tracts were analyzed: uncinate fasciculus (UF), inferior longitudinal fasciculus (ILF), inferior fronto-occipital fasciculus (IFOF), cingulum, and corticospinal tract (CST; see Figure 1). As a final step, for each fiber pathway of interest, for each hemisphere, and for each subject, FA, RD, and AD statistics were exported and further analyzed.

### Brain-Behavior Data Analyses

Analyses were conducted using R v.3.5.3 (Team, 2020). As an initial step, data were inspected for missingness. Only 2% of all data were missing, and thus multiple imputation across 20 imputations (using R package Multivariate Imputation by Chained Equations; *mice*) was performed. We also examined whether there were significant group differences when it came to movement in the scanner. Out of 102 directions, the ADHD sample moved more frequently and lost more directions (*M*= 83.96 directions kept, *SD*= 12.53) compared to TD (*M*= 88.74, *SD=* 9.29; (*t*(196)= 3.03, *p* = 0.0027). We included the number of retained diffusion directions as a covariate in all subsequent models. In these regression models, we used robust regression (R function *rlm*; (Wright & London, 2009)) given that they are less influenced by outlying values (Wilcox, 2012). We also improved the estimation of the reliability of the parameter estimate by using the bootstrap method (Efron, 1981, 1987) to calculate the standard errors and 95% confidence intervals.

## Results

First, we examined TD vs. ADHD group differences in behavioral measures, and in the diffusion metrics across the five fiber tracts (UF, IFOF, Cingulum, ILF, and CST). As expected, children in the ADHD sample had significantly higher rates of CP (*t(196)* = 11.78*, d* = 1.69, *p* < 0.0001) and CU behaviors (*t(196)* = 5.60*, d* = 0.79, *p* < 0.0001) compared to TD children. There was also a significant group difference on IQ (*t(194) = −4.31*, *d* = −0.62, *p* < 0.0001), with the ADHD group (*M* = 96.24, *SD* = 1.29) scoring lower than the TD group (*M* = 103.67, *SD* = 1.13).

Because FA is calculated using information contained in the other metrics (e.g., RD and AD), and because it is the most commonly reported summary metric for DWI, we focus on FA differences in our results. We found no significant group differences for whole brain FA (*t*(196) = 0.79, d = 0.11, p = 0.43), nor for bilateral cingulum (*t*(196) = 0.25, *d* = 0.04, *p* = 0.80 for left; *t*(196) = 0.002, d = 0.0., *p* = 0.99 for right). However, all other pathway differences for FA were statistically significant (all *p* < .0001; see Figure 2).

**Figure 2.**
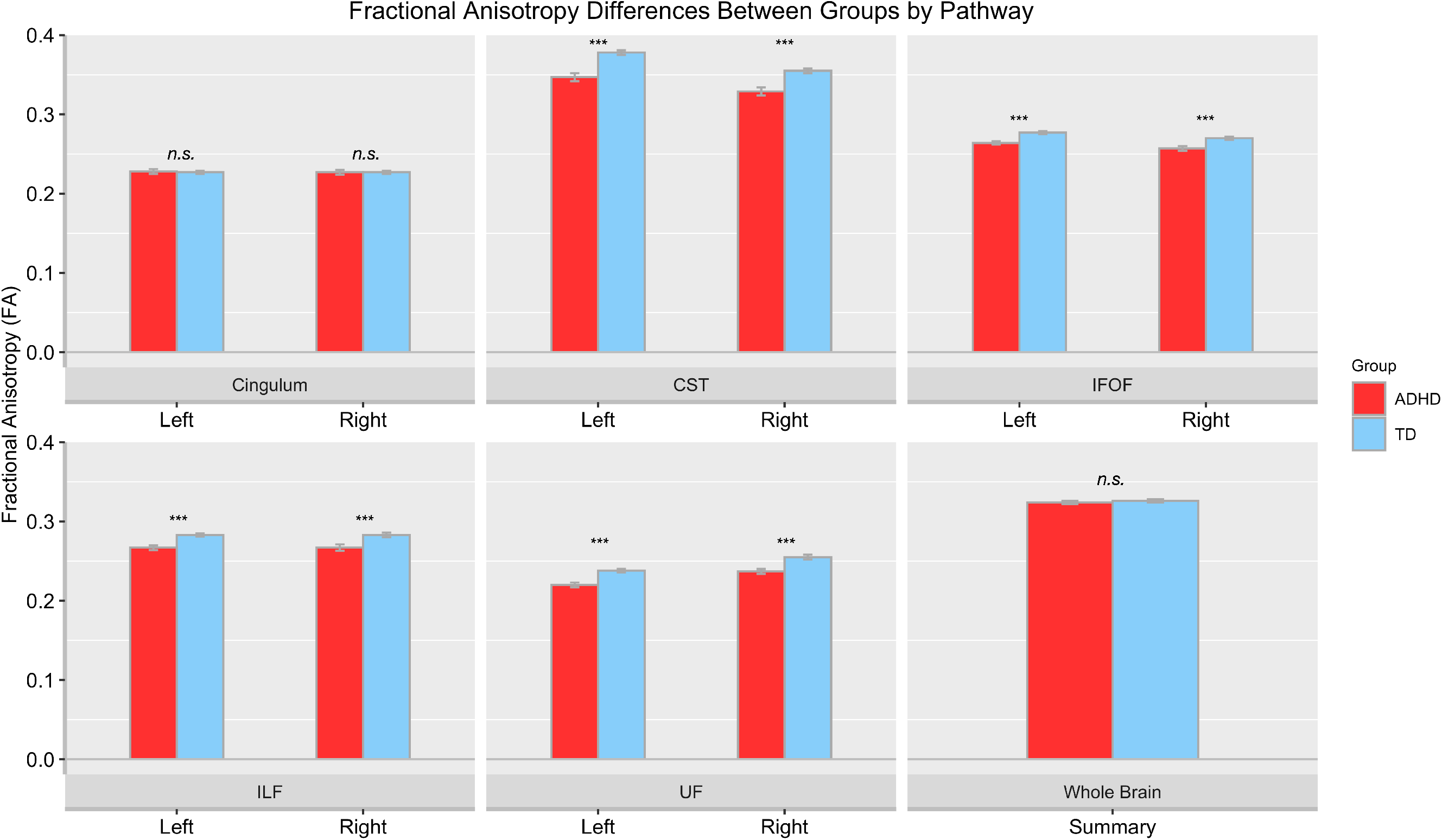
Fractional anisotropy (FA) mean differences are plotted by group, for each pathway and for each hemisphere. TD = typically developing children, ADHD = Attention-Deficit/Hyperactivity Disorder, CST = Corticospinal Tract; IFOF = Inferior fronto-occipital fasciculus; ILF = Inferior longitudinal fasciculus; UF = Uncinate fasciculus. The average of the whole-brain FA is also plotted. *n.s.* = non-significant. *** *p* < .001.

The next two analyses focused on the ADHD group, who as noted above showed substantially higher rates of CP and CU behaviors. In the second analysis, using robust regression we examined whether any of the examined fiber pathways were associated with CP. Again, we focus on FA, but to be comprehensive these models were run for AD and RD diffusion metrics as well. These analyses controlled for sex, whole-brain diffusion (e.g., for FA, AD, and RD we controlled for whole-brain FA, AD, and RD respectively), movement, SES, IQ, hyperactivity/impulsivity, and inattention. Results are reported in Table 1, and show no significant associations between FA of any of the fiber pathways and CP symptoms.

**Table 1.** Results of robust regressions examining the association between tract diffusivity metric and conduct problems (CP).

| Tract -> Conduct Problems (CP) | *B* (*SE*) | $\beta$ | *t* | *p* | CI 95% |
| --- | --- | --- | --- | --- | --- |
| **Left Hemisphere** |  |  |  |  |  |
| UF FA | -0.95 (1.39) | -0.06 | -0.68 | 0.50 | -3.66, 1.77 |
| Cingulum FA | 0.09 (1.77) | 0.01 | 0.05 | 0.96 | -3.39, 3.56 |
| ILF FA | -0.89 (1.67) | -0.05 | -0.53 | 0.59 | -4.16, 2.38 |
| IFOF FA | -2.59 (1.87) | -0.13 | -1.39 | 0.17 | -6.24, 1.07 |
| CST FA | 0.09 (0.90) | 0.01 | 0.10 | 0.92 | -1.67, 1.84 |
| UF AD | -1.03 (2.13) | -0.05 | -0.49 | 0.63 | -5.20, 3.14 |
| Cingulum AD | 0.69 (1.86) | 0.04 | 0.37 | 0.71 | -2.96, 4.34 |
| ILF AD | -0.58 (1.52) | -0.04 | -0.38 | 0.70 | -3.56, 2.40 |
| IFOF AD | 0.83 (2.38) | 0.04 | 0.35 | 0.73 | -3.83, 5.49 |
| CST AD | 0.33 (2.58) | 0.01 | 0.13 | 0.90 | -4.74, 5.39 |
| UF RD | 0.15 (1.42) | 0.01 | 0.11 | 0.92 | -2.64, 2.94 |
| Cingulum RD | -1.62 (2.35) | -0.09 | -0.69 | 0.49 | -6.23, 3.00 |
| ILF RD | -1.16 (2.81) | -0.06 | -0.41 | 0.68 | -6.67, 4.36 |
| IFOF RD | 2.87 (2.11) | 0.14 | 1.30 | 0.20 | -1.47, 7.20 |
| CST RD | -0.16 (1.00) | -0.01 | -0.16 | 0.87 | -2.12, 1.80 |
| **Right Hemisphere** |  |  |  |  |  |
| UF FA | -0.77 (1.36) | -0.05 | -0.56 | 0.57 | -3.44, 1.91 |
| Cingulum FA | 0.48 (1.81) | 0.03 | 0.27 | 0.79 | -3.07, 4.04 |
| ILF FA | 0.38 (1.26) | 0.03 | 0.30 | 0.76 | -2.09, 2.85 |
| IFOF FA | -2.54 (1.74) | -0.14 | -1.46 | 0.15 | -5.95, 0.87 |
| CST FA | 0.47 (0.94) | 0.05 | 0.50 | 0.62 | -1.37, 2.30 |
| UF AD | 1.11 (2.55) | 0.04 | 0.43 | 0.67 | -3.89, 6.10 |
| Cingulum AD | 0.30 (1.83) | 0.02 | 0.16 | 0.87 | -3.29, 3.89 |
| ILF AD | 1.08 (1.72) | 0.06 | 0.63 | 0.53 | -2.29, 4.44 |
| IFOF AD | -1.15 (2.49) | -0.05 | -0.46 | 0.65 | -6.04, 3.74 |
| CST AD | 3.17 (2.48) | 0.13 | 1.28 | 0.21 | -1.70, 8.04 |
| UF RD | 0.55 (1.43) | 0.04 | 0.39 | 0.70 | -2.25, 3.36 |
| Cingulum RD | -2.50 (2.38) | -0.12 | -1.05 | 0.30 | -7.17, 2.17 |
| ILF RD | -0.42 (1.89) | -0.02 | -0.22 | 0.82 | -4.12, 3.27 |
| IFOF RD | 2.03 (2.34) | 0.10 | 0.87 | 0.39 | -2.55, 6.61 |
| CST RD | -0.30 (0.97) | -0.03 | -0.31 | 0.76 | -2.19, 1.60 |

Note. All regressions controlled for the following: sex, whole brain diffusion (either FA, AD, or RD depending on predictor of interest), movement in the scanner, socioeconomic status, IQ, hyperactivity/impulsivity, and inattention. UF = Uncinate fasciculus; ILF = Inferior longitudinal fasciculus; IFOF = Inferior fronto-occipital fasciculus; CST = Corticospinal tract; CI = Confidence Interval. FA = Fractional Anisotropy; AD = Axial Diffusivity; RD = Radial Diffusivity. \* $p < .05$ .

In the third analysis, the same models were run, but the ICU composite measuring CU behaviors was substituted for the outcome variable, and CP symptoms were entered as an additional covariate. Table 2 shows that, for FA, bilateral UF and left IFOF FA are negatively associated with CU behaviors. Consistent with these findings, bilateral AD was negatively associated with CU behaviors, which suggests that the finding for FA is driven mainly by the longitudinal component of the diffusion tensor. Taken together, these results show that, even when controlling for whole brain diffusion differences, demographic effects, ADHD symptom severity, and CP, reduced directional diffusion within bilateral UF and left IFOF fiber pathways is significantly associated with increased CU behaviors. Figure 3 shows these effects plotted for the left and right UF, and left IFOF.

**Table 2.** Results of robust regressions examining the association between tract diffusivity metric & callous-unemotional (CU) behaviors.

| Tract -> Callous-Unemotional (CU) Behaviors | *B* (*SE*) | $\beta$ | *t* | *p* | CI 95% |
| --- | --- | --- | --- | --- | --- |
| **Left Hemisphere** |  |  |  |  |  |
| UF FA | -2.94 (-1.13) | -0.18 | -2.27 | **0.026 \*** | -5.48, -0.40 |
| Cingulum FA | 2.18 (-1.78) | 0.12 | 1.23 | 0.22 | -1.31, 5.67 |
| ILF FA | -2.21 (-1.64) | -0.11 | -1.30 | 0.20 | -5.34, -0.77 |
| IFOF FA | -4.25 (-1.78) | -0.20 | -2.40 | **0.018 \*** | -7.73, -0.77 |
| CST FA | -0.37 (-0.09) | -0.04 | -0.40 | 0.69 | -2.16, 1.42 |
| UF AD | -4.12 (-2.03) | -0.17 | -2.03 | **0.045 \*** | -8.10, -0.14 |
| Cingulum AD | 2.17 (-1.89) | 0.11 | 1.15 | 0.25 | -1.54, 5.87 |
| ILF AD | -2.77 (-1.41) | -0.17 | -1.97 | 0.05 | -5.53, -0.02 |
| IFOF AD | -3.92 (-2.31) | -0.15 | -1.69 | 0.09 | -8.45, 0.62 |
| CST AD | 0.09 (-2.63) | 0.00 | 0.04 | 0.97 | -5.07, 5.26 |
| UF RD | 1.91 (-1.43) | 0.12 | 1.34 | 0.18 | -0.89, 4.70 |
| Cingulum RD | -1.76 (-2.45) | -0.09 | -0.72 | 0.47 | -6.56, 3.04 |
| ILF RD | -0.94 (-2.81) | -0.04 | -0.34 | 0.74 | -6.46, 4.57 |
| IFOF RD | 3.16 (-2.22) | 0.15 | 1.43 | 0.16 | -1.18, 7.51 |
| CST RD | 0.48 (-1.04) | 0.04 | 0.46 | 0.64 | -1.56, 2.52 |
| **Right Hemisphere** |  |  |  |  |  |
| UF FA | -2.70 (-1.30) | -0.17 | -2.08 | **0.040 \*** | -5.25, -0.15 |
| Cingulum FA | 0.86 (-1.88) | 0.05 | 0.46 | 0.65 | -2.82, 4.55 |
| ILF FA | -1.51 (-1.24) | -0.11 | -1.22 | 0.23 | -3.95, 0.92 |
| IFOF FA | -2.49 (-1.75) | -0.13 | -1.43 | 0.16 | -5.91, 0.94 |
| CST FA | -0.26 (-0.94) | -0.28 | -0.28 | 0.78 | -2.10, 1.58 |
| UF AD | -6.83 (-2.26) | -0.24 | -3.03 | **0.003 \*\*** | -11.26, -2.41 |
| Cingulum AD | 2.30 (-1.85) | 0.12 | 1.24 | 0.22 | -1.33, 5.92 |
| ILF AD | -1.54 (-1.72) | -0.09 | -0.90 | 0.37 | -4.91, 1.82 |
| IFOF AD | -0.35 (-2.55) | -0.01 | -0.14 | 0.89 | -5.35, 4.65 |
| CST AD | -1.71 (-2.58) | -0.07 | -0.66 | 0.51 | -6.76, 3.34 |
| UF RD | 1.31 (-1.44) | 0.08 | 0.90 | 0.37 | -1.53, 4.13 |
| Cingulum RD | 0.66 (-2.44) | 0.03 | 0.27 | 0.79 | -4.13, 5.44 |
| ILF RD | 1.52 (-1.94) | 0.08 | 0.78 | 0.44 | -2.28, 5.31 |
| IFOF RD | 3.37 (-2.21) | 0.16 | 1.52 | 0.13 | -0.96, 7.70 |
| CST RD | 0.27 (-1.00) | 0.02 | 0.27 | 0.79 | -1.69, 2.24 |

Note. All regressions controlled for the following: sex, whole brain diffusion (either FA, AD, or RD depending on predictor of interest), movement in the scanner, socioeconomic status, intelligence, hyperactivity/impulsivity, inattention, and conduct problems (CP). UF = Uncinate fasciculus; ILF = Inferior longitudinal fasciculus; IFOF = Inferior fronto-occipital fasciculus; CST = Corticospinal tract; CI = Confidence Interval. FA = Fractional Anisotropy; AD = Axial Diffusivity; RD = Radial Diffusivity. \* $p < .05$ . \*\* $p < .01$ .

**Figure 3.**
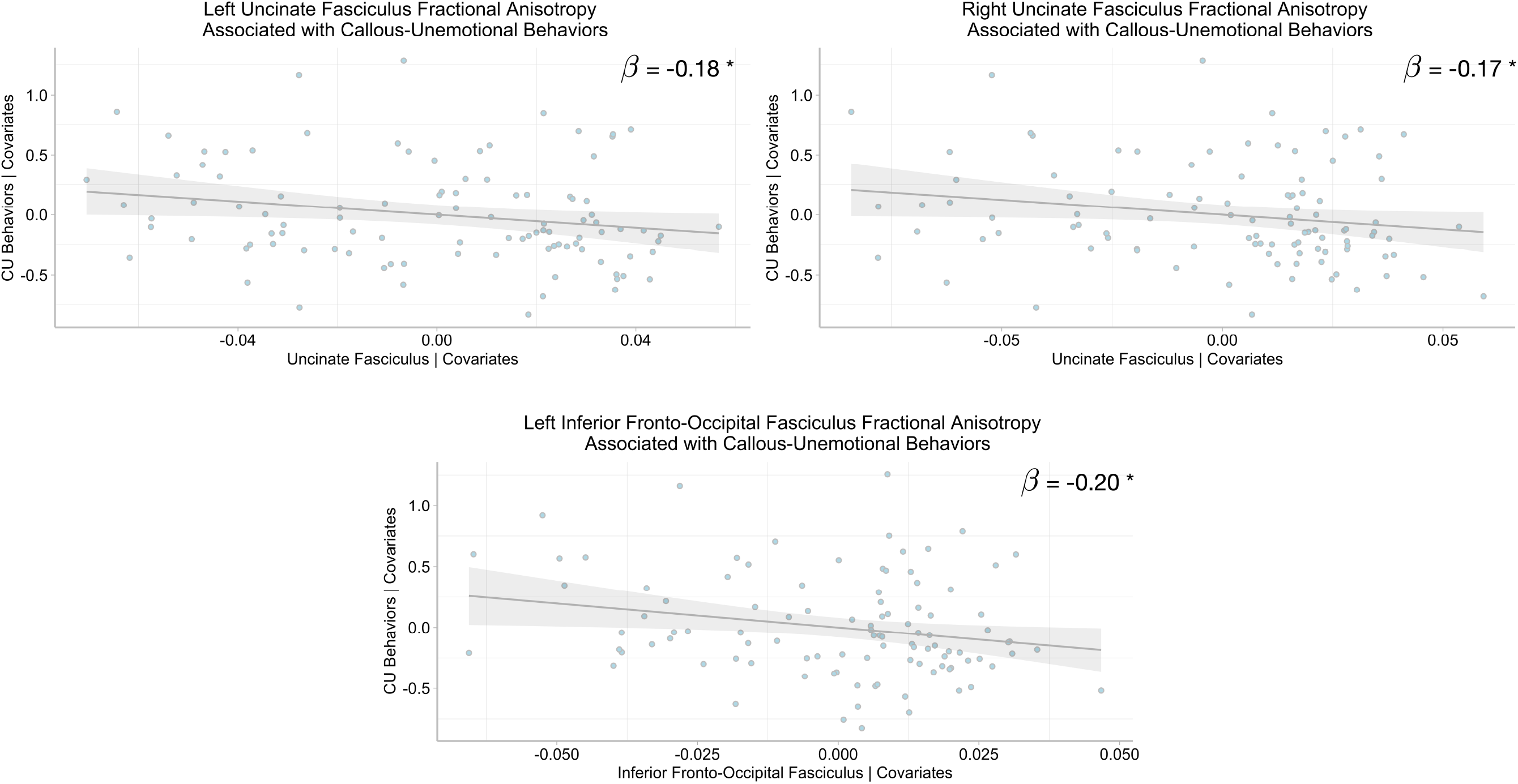
Added-variable plots show associations between bilateral uncinate fasciculus and left inferior fronto-occipital fasciculus fractional anisotropy (FA) and callous-unemotional (CU) behaviors, controlling for the following covariates: sex, whole brain FA, movement in the scanner, socioeconomic status, intelligence, hyperactivity/impulsivity, inattention, and conduct problems (CP). β= standardized regression slope parameter. Shading = 95% confidence intervals. * *p* < 0.05.

## Discussion

In a recent review, Waller and colleagues (2017) found that reduced FA in the fiber tracks connecting the extended limbic, frontal, and temporal regions (namely UF, cingulum, IFOF, and ILF tracks investigated here) is associated with antisocial behavior in adults. In adolescents, they showed that the results were more mixed, with some studies showing reduced FA in these tracks, and others showing increased FA. In our study of younger children, the first to our knowledge with a large predominantly pre-kindergarten sample (M^age^= 5.66), we found that relative to TD children, children with ADHD had reduced FA across the IFOF, ILF, UF, and CST. Of note, although the CST was included as a control tract, previous studies have documented that children with ADHD have reduced FA in this pathway as well, relative to TD children (D’Agati, Casarelli, Pitzianti, & Pasini, 2010; Hamilton et al., 2008). This reduced integrity of the CST may be associated with fine and gross motor difficulties, consistent hallmarks of ADHD (Mokobane, Pillay, & Meyer, 2019). Thus, our results replicate, in younger children, well-established findings regarding the group differences between youth diagnosed with ADHD and TD youth (e.g., see (Svatkova et al., 2016; Wu et al., 2020).

Notably, we did not find a difference in general FA across the whole brain, nor did we find a group difference in the cingulum. Differences in FA in cingulum have been found previously in studies of older children with ADHD (Svatkova et al., 2016; Wu et al., 2020), but not in all such studies (Ashtari et al., 2005; Davenport, Karatekin, White, & Lim, 2010), and not in children with CU behaviors (Pape et al., 2015) or CD (Finger et al., 2012; Haney-Caron et al., 2014). It is important to point out, though, that these studies showing differences have tended to be small sample studies (e.g., *n* < 30), increasing the possibility that such effects are spurious ((2015) is a notable exception). Our study is a comparatively large sample, and thus it is reasonable to conclude that, at this age, cingulum FA is not different between ADHD and TD groups. We interpret this as an important potential null finding as it relates to understanding the broader circuit dysfunction in youth with ADHD. However, it is possible differences might arise later in development, as fiber pathways continue to show maturational change well beyond the pre-school and early school-age period that we studied here.

Within the ADHD group analyses, while we found no associations with any white matter properties and CP, we did find reliable associations with CU behaviors. Turning first to the null findings for CP, there are a number of obvious methodological differences with our study and prior studies. First, we examined a very young group of children, and previous work has mostly focused on adolescents. Second, within the clinical group, all children in our study were diagnosed with ADHD, which may contribute to the mixed pattern of findings in the literature (2017). More work is clearly needed examining the white matter microstructure within the frontal-temporal-extended limbic system taking into account ADHD comorbidity.

Our most noteworthy finding within the ADHD group is that even after accounting for ADHD symptom severity, CP, demographic variables, and whole brain FA, CU behaviors were independently related to reduced FA in bilateral UF and left IFOF. Both of these fiber pathways support connections of temporal lobe and limbic structures with orbitofrontal cortex, and both pathways have been associated with CU behaviors in adolescents (e.g., (Breeden et al., 2015; Pape et al., 2015; Sarkar et al., 2013) and psychopathy in adults (see Waller et al., (2017) for review). Both pathways have also been associated with emotion regulation. The UF supports extensive connectivity with amygdala and orbitofrontal cortex, and not surprisingly it has been implicated in the recognition of facial expressions of emotions (Philippi, Mehta, Grabowski, Adolphs, & Rudrauf, 2009). Indeed, emotion processing deficits that include not only reduced amygdala response to fearful faces, but also general emotion recognition deficits at the behavioral level, have been consistently associated with CU behaviors (Dadds, Kimonis, Schollar-Root, Moul, & Hawes, 2018; Marsh et al., 2008). The IFOF is a far more extensive fiber pathway that passes just dorsal to the UF in its anterior course, but it also supports emotion recognition (Unger, Alm, Collins, O’Leary, & Olson, 2016). Disrupted emotional responsiveness is potentially a core feature in at least a subset of children displaying CU behaviors (Frick & Viding, 2009; Northam & Dadds, 2020). Our results suggest that disruption of the main fiber pathways supporting emotional processing might be a contributing factor to the development of CU behaviors in such children. Further, these differences can be detected reasonably early in development (i.e., in the preschool/early school-age period). The results thus add an additional level of analysis on which to advance causal theories for the development of CU behaviors (Frick & Viding, 2009).

It is important to note that the findings with respect to bilateral UF were mainly driven by the longitudinal component of the diffusion tensor (i.e., AD). While speculative, as we do not have access to the specific microstructural properties of the brain tissue, it is the case that AD is more sensitive to disruptions of axonal integrity and packing density, while RD quantifications are more sensitive myelin integrity (Basser et al., 1994; Winston, 2012). This may suggest that the UF of children with CU behaviors is characterized by less coherent longitudinal fiber orientation rather than reduced or delayed myelination, although such a possibility would need additional verification. Regardless, our findings add to the extant pediatric literature highlighting the importance of the disrupted connectivity between amygdala and orbitofrontal cortex as it relates specifically to CU traits/behaviors (Blair, 2007).

Some limitations to the current study include not having a pure CP group as our clinical sample had a primary diagnosis of ADHD. Given the high comorbidity of CP and ADHD in young children (Bendiksen et al., 2017), our approach was to isolate the CP component by statistically controlling for ADHD severity. However, we acknowledge the limitations of statistically covarying versus obtaining a pure CP group, and thus we cannot rule out the fact that the widespread disruption of multiple fiber tracts that we found in our ADHD sample may be similar within a “pure” CP sample. Second, while we focused on several major fiber tracts related to network of extended limbic, frontal, and temporal regions given their theoretical and empirical associations with the development of CP/CU and associated impairments, it will be important in the future to also examine the fronto-striatal-cerebellar neurocircuitry given its link to ADHD (van Ewijk, Heslenfeld, Zwiers, Buitelaar, & Oosterlaan, 2012). Lastly, another limitation of the current study is the homogeneity of the sample, which was largely Latinx (80%) due to the study’s geographical location. The homogeneity of the sample limits the generalizability of these findings but can be viewed as a strength as Latinx children represent the fastest growing group in the U.S., but are understudied in child psychopathology research (La Greca, Silverman, & Lochman, 2009).

In sum, relative to TD children, children with ADHD (with high comorbidity for CP) were found to have white matter disruption on four out of the five fiber tracks we examined (except for cingulum). Within the ADHD group, we did not find any associations between CP and reduced white matter integrity. However, we did find that CU behaviors were associated with reduced FA in bilateral UF and left IFOF. Consistent with the adult and limited adolescent literature, our results suggest that these pathways may be biomarkers of CU behaviors/traits even in very young children. Such white matter disruptions within the frontal/limbic network map onto the emotional processing deficits, including lack of empathy, that are the core features of CU behaviors. Moving forward it will be important to identify multiple biomarkers (i.e., a “biosignature”) which may help guide us into the developing more targeted treatments and individualizing treatment options for young children with CP and who display elevated levels of CU behaviors.

## Acknowledgements

We would like to acknowledge the support of Miami-Dade County Public Schools and thank the families and dedicated staff who participated in the study.

## Footnotes

* Footnote: given the young age of our sample and to facilitate consistency in our terminology when reviewing the literature, we used the term CU *behaviors* throughout the paper although we acknowledge that in older samples the term CU *traits* is also frequently used.

